# csuWGCNA: a combination of signed and unsigned WGCNA to capture negative correlations

**DOI:** 10.1101/288225

**Authors:** Rujia Dai, Yan Xia, Chunyu Liu, Chao Chen

## Abstract

Network analysis helps us to understand how genes jointly affect biological functions. Weighted Gene Co-expression Network Analysis (WGCNA) is a frequently used method to build gene co-expression networks. WGCNA may be calculated with signed or unsigned correlations, with both methods having strengths and weaknesses, but both methods fail to capture weak and moderate negative correlations, which may be important in gene regulation. Combining the advantages and removing the disadvantages of both methods in one analysis would be desirable. In this study, we present a combination of signed and unsigned WGCNA (csuWGCNA), which combines the signed and unsigned methods and improves the detection of negative correlations. We applied csuWGCNA in 14 simulated datasets, six ground truth datasets and two large human brain datasets. Multiple metrics were used to evaluate csuWGCNA at gene pair and gene module levels. We found that csuWGCNA provides robust module detection and captures more negative correlations than the other methods, and is especially useful for non-coding RNA such as microRNA (miRNA) and long non-coding RNA (lncRNA). csuWGCNA enables detection of more informative modules with biological functions than signed or unsigned WGCNA, which enables discovery of novel gene regulation and helps interpretations in systems biology.

## Introduction

Biological functions are controlled by a group of co-regulated or co-expressed genes in the context of a network in systems biology. Network analyses helps to explore system-level functionality of genes, such as global interpretation of transcriptome and putative regulation relationships between genes. Numerous approaches and algorithms have been proposed for module detection in gene expression data^1^.

Weighted Gene Co-expression Network Analysis (WGCNA) is a widely used method for detecting important gene pairs and modules^2^. WGCNA is completed in three steps, the first being construction of gene co-expression networks (GCNs) from a matrix of the correlations between expression of genes across samples. Then an adjacency matrix is constructed based on the correlation matrix. The genes and their connection strengths are regarded as nodes and edges, respectively, in the GCNs. In the second step, gene modules are obtained via hierarchical clustering and tree cutting. Lastly gene modules are related to external information to interpret their biological functions and gene regulation in modules can also be revealed. Detecting the group or module of co-expressed genes has generated insights in brain transcriptome architecture^3^, evolution^4^, aging^5^, cell diversity^5^, and psychiatric disorders^6,7^.

The adjacency matrix is the foundation of the WGCNA procedure. According to the definition of adjacency matrix, WGCNA can be classified into two major types: signed method and unsigned method. The two methods treat negative correlations differently. Consider the gene expression matrix G_mxn_, where m is the number of genes and n is the number of samples. The WGCNA procedure generates a correlation matrix S from G via pair-wise correlations^8^. Then the adjacency matrix A is constructed from S, depending on whether the adjacency is signed or unsigned. In the signed method, positive correlations are prioritized over the negative correlations. Larger positive correlations have larger adjacency, and larger negative correlations have smaller adjacency. The adjacency of strong negative correlations is close to zero. In the unsigned method, negative and positive correlations are considered equally. Adjacency is only determined by the size of correlations without considering direction. The adjacency of strong negative correlations is close to 1. Adjacency for genes i and j is defined as follows in these two methods, where power β is set to keep the network with scale-free topology property^9^ (Only a few nodes in the network are highly connected and most of the gene are connected with a few genes).

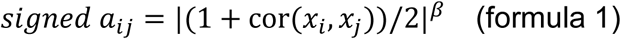

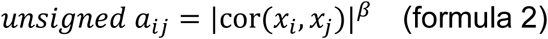

The signed and unsigned methods have advantages and disadvantages. The modules detected by the signed method are more robust to their biological functions than unsigned modules. A previous study on embryonic stem cells showed that the signed method identifies modules with more specific expression patterns than the unsigned method^10^. The unsigned method can capture more negatively correlated genes than the signed method, for example, non-coding RNA and their targets. Detecting this type of negatively correlated genes is difficult because they are considered not connected by the signed method. However, the unsigned method is only capable of detecting strong correlations. Negative regulation relationships in biological systems are usually weak or moderate so they will not be detected using the unsigned method. For example, a study reported that the more than 50% of microRNA (miRNA)-mRNA correlations in 35 human tissues were 0~-0.3 and only one correlation was lower than −0.5^11^.

Regulation by suppression is common in functional biology pathways. For example, miRNA and long non-coding RNA (lncRNA) are two types of non-coding RNAs reported to repress target genes^12–14^. MiRNAs can regulate gene transcription and inhibit translation of mRNA^15–17^. Brain-specific miRNA miR-134 was reported to inhibit Limk1 translation in mice and may contribute to synaptic development^18^. LncRNA is another type of regulating RNA, and 40% of the known lncRNAs are expressed specfically in brain^19^. LncRNAs have been implicated in regulating gene expression at diverse levels, such as transcription, RNA processing and translation^20–21^. In the nucleus, lncRNA regulates the gene by interacting with chromatin-modifying complex or transcriptional factors. For example, the lncRNA RMST has been reported to be down-regulated by the transcriptional factor REST, and RMST regulates neurogenesis by binding SOX2 in vitro^22^. LncRNA BDNF-AS is the natural antisense transcript to BDNF, itself a key contributor to synaptic function^23^. By dynamically repressing BDNF expression in response to neuronal depolarization, BDNF-AS modulates synaptic function. Researchers should be careful not to neglect moderate repression roles in co-expression networks. Both miRNA and lncRNA are likely to have negative correlations with other genes and are likely to be undetected by WGCNA.

In this study, we developed a new method named csuWGCNA that combines signed and unsigned WGCNA methods to detect more negative correlations. We used 14 simulation data sets, 8 ground truth data sets with known modules, and two brain gene expression data sets from Stanley Medical Research Institute and the PsychENCODE project to comprehensively evaluate signed, unsigned, and csuWGCNA methods at pairwise gene and module levels. We showed that csuWGCNA is more effective at capturing negative correlations such as those involving miRNA-target and lncRNA-gene pairs. We also showed that csuWGCNA can robustly detect modules with biological functions. This method balances the signed and unsigned methods and provides a more effective way to analyze whole-transcriptome data.

## Results

### Combining signed and unsigned WGCNA

To combine the signed and unsigned WGCNA, we developed a new method named csuWGCNA for gene co-expression network construction. The csuWGCNA defines the adjacency matrix as follows,

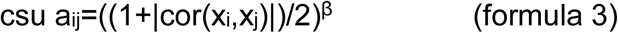

For the genes i, and j, cor(x_i_,x_j_) is the correlation and a_ij_ is the adjacency between them. This method combines adjacency calculations used in signed and unsigned methods (Figure 1A) and csuWGCNA considers weak, moderate, and strong correlations as well as their direction (Figure 1B). The process of csuWGCNA includes power selection, adjacency calculation based on similarity matrix, topological overlap Matrix (TOM) construction, hierarchical clustering, dynamic tree cutting, and module merging.

**Figure 1.**
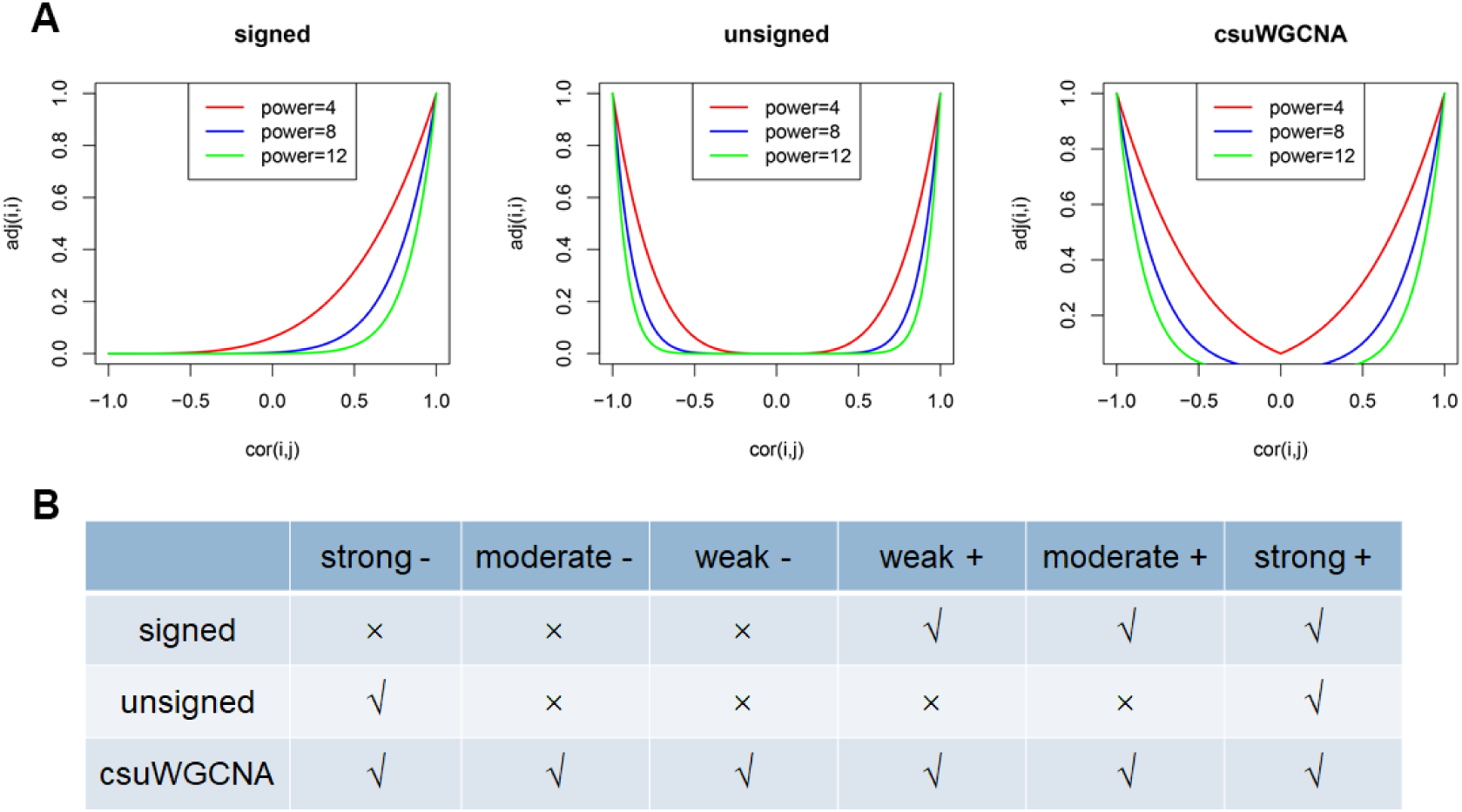
The correlations captured by three methods for gene co-expression network construction. (A). Network adjacency (y-axis) versus correlation (x-axis) for weighted networks for the signed method, unsigned method, and csuWGCNA. The color of the line denotes the power used. Note that correlation=-1 leads to adjacency = 0 in the signed network and adjacency =1 in the unsigned and csuWGCNA network. (B). The types of correlations captured by three networks. The strong and weak denote the degree of correlations, the “-” and “+” denote the direction of correlations. The “√” and “×” denote the possibility of capturing the corresponding types of correlation.

### Evaluation workflow

We evaluated signed, unsigned, and csuWGCNA methods on 22 gene expression datasets (Figure 2). We tested 14 simulated datasets, six ground truth datasets with known regulation networks for *E. coli*, yeast and synthetic data, and two real gene expression datasets from human brains (Figure 2A). We set three types of metrics to evaluate the three methods at the gene pair and gene module levels. To evaluate the capture of negative correlations, we examined the proportion of negatively correlated gene pairs in observed modules and their targets. To compare the observed modules with known modules, we chose six metrics: specificity, sensitivity, positive predictive value (PPV), negative predictive value (NPV), recovery, and relevance. To evaluate the biological functions of observed modules, we applied false discovery rate (FDR) to predict the enrichment of Kyoto Encyclopedia of Genes and Genomes (KEGG) pathway. The final score for each method was the sum of z-scores of all metrics used.

**Figure 2.**
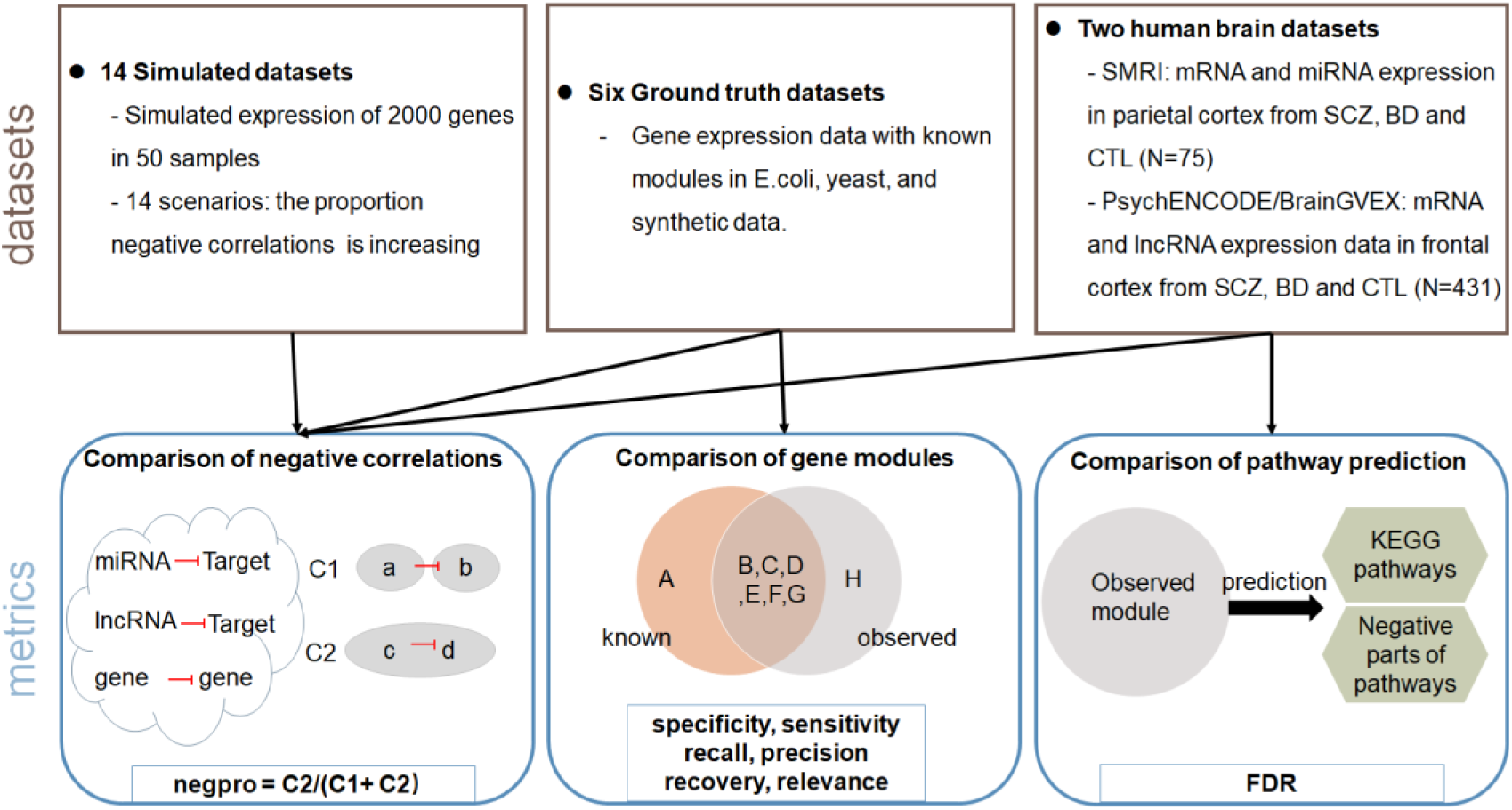
Overview of evaluation. Up panel is total 22 datasets used in this evaluation which including simulation data, ground truth data and human brain data. Bottom panel is the metrics used to evaluate the performance of the method which classified in three categories: detection negative correlations, comparison with known modules, and biological pathway prediction. The arrow indicates the combination of datasets and metrics.

### csuWGCNA captures more weak-negative correlations than signed and unsigned methods in simulation analysis

To evaluate the detection of negative correlations, we simulated gene expression of 2000 genes from 50 samples in 14 scenarios with increased proportions of negative correlations. To assess the types of correlations captured, we classified all correlations as follows: weak (|bicor|<0.3), moderate (0.3≤|bicor|≤0.6), and strong (|bicor|>0.6). Based on the predicted adjacency curves for the three methods, the csuWGCNA was expected to detect more negative correlations than the unsigned method, especially for weak and moderate correlations (Figure 3A).

**Figure 3.**
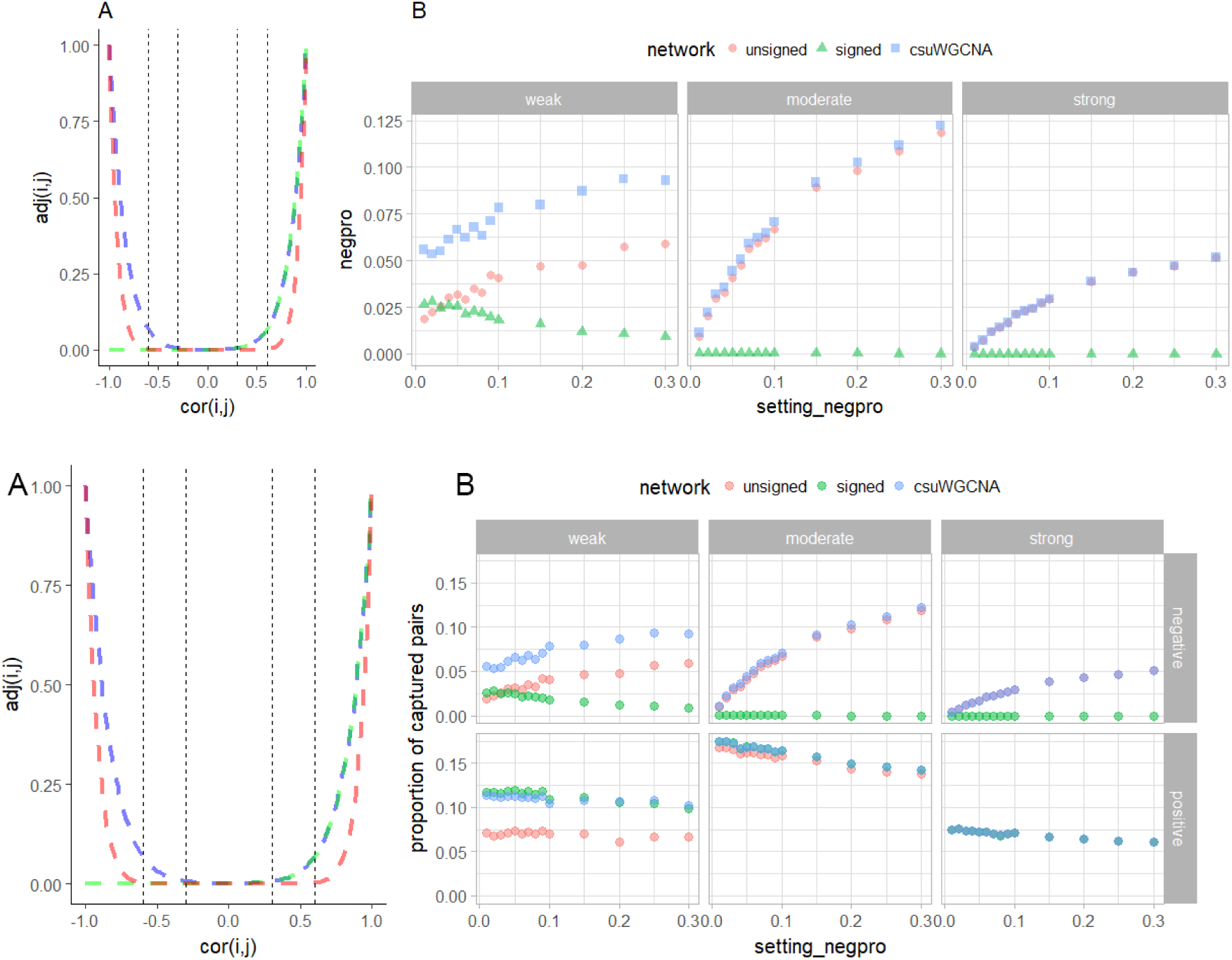
simulation result. (A) distribution of adjacency calculated by three methods for gene co-expression network construction. The dashed line denotes the boundary of weak, moderate and strong correlations. (B) the proportion of negative correlation captured by the three methods. The axis is the proportion of negative correlations pre-set in each simulation data.

The simulation result showed that csuWGCNA captured 5%~9% of the weak correlations which is higher than that of both of unsigned and signed methods, respectively (Bonferroni-Holm (BH) adjusted P_csuWGCNA-signed_ =5.2e-05, P_csuWGCNA-unsigned_ =1.1e-04, two-sample Wilcoxon test). csuWGCNA captured 1%~12% of the moderate correlations, which is significantly higher than that of signed method (BH adjusted P =5.2e-05) but only 5% higher than that of unsigned methods with an insignificant p-value 0.73. Both csuWGCNA and unsigned method capture 0.4%~5% strong correlations while the signed method is unable to detect strong correlations (BH adjusted P =1.5e-05 for both signed-csuWGCNA and signed-unsigned comparison). Meanwhile, csuWGCNA and the unsigned method captured an increasing proportion of negative correlations when the proportion of negative correlations in the data increased. In contrast, the performance of the signed method did not change with increasing numbers of negative correlations and had poor detection throughout the range. The simulation result was in accord with the adjacency distribution and suggests that csuWGCNA is capable of capturing more negatively correlated gene pairs than both signed and unsigned methods, especially for those that are weakly correlated.

### csuWGCNA captures more negative correlations and retains robust module reproducibility in ground truth data

To evaluate the three methods at the gene module level, we used six ground truth datasets with gene expression and corresponding known modules for *E. coli*, yeast and synthetic regulatory networks.

We evaluated module reproducibility using the following metrics: positive predictive value (PPV), negative predictive value (NPV), recovery, relevance, sensitivity and specificity (Supplemental Figure). We synthesized the six metrics into one metric named “module score” to represent module reproducibility. We found that the module score of csuWGCNA is slightly higher than that of signed and unsigned methods with insignificant p-value (Figure 4A).

We considered negative correlations in this ground truth comparison. The proportion of negative correlations captured by gene modules (negpros) of csuWGCNA and the unsigned methods were higher than that of the signed method (BH adjusted P_csuWGCNA-signed_ =0.006, P_csuWGCNA-unsigned_ =0.015, two-sample Wilcoxon tests, Figure 4B). No difference was detected between negpro of csuWGCNA and unsigned method.

**Figure 4.**
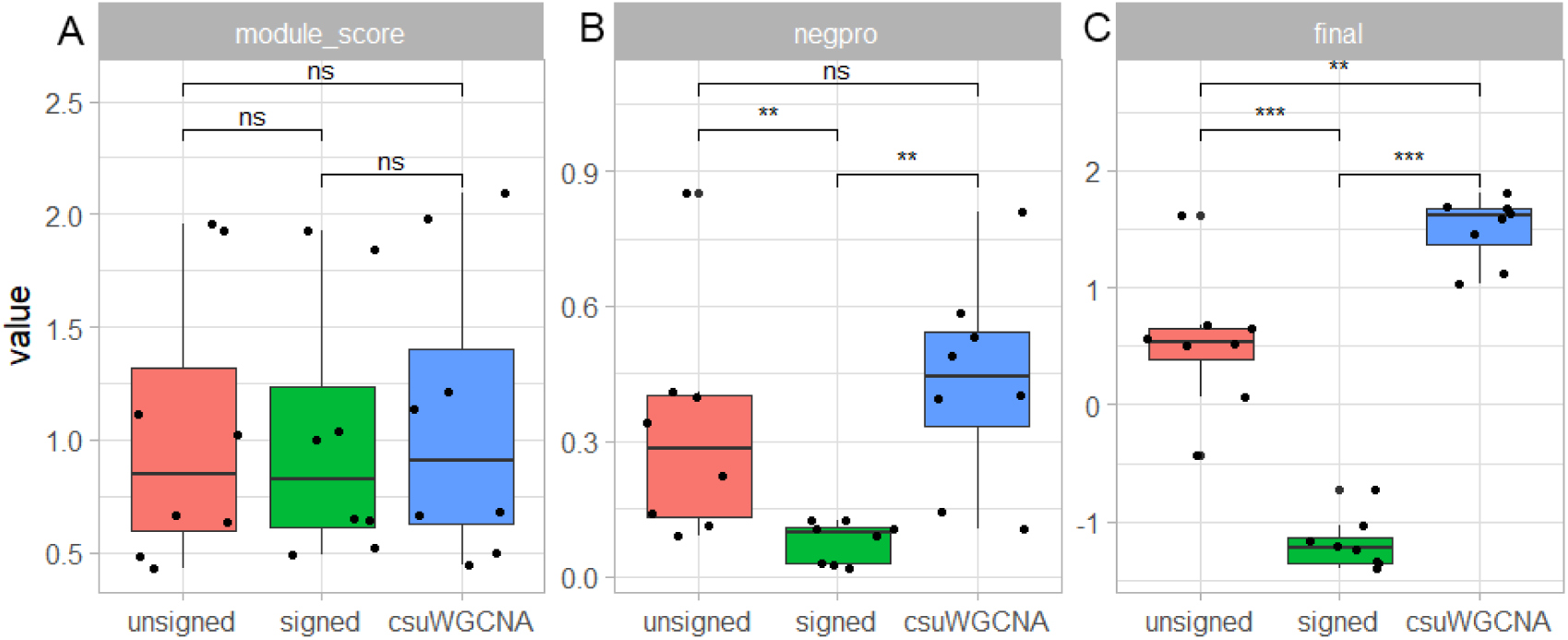
Overall performance of signed, unsigned and csuWGCNA in ground truth data. (A) The module score synthesizing individual metrics. (B) The negpro of three methods in given dataset. (C) The final score synthesizing module score and negpro. Two-sample Wilcoxon test was used to test the difference (N=8). ns denotes non-significance. “**” and “***” denotes p-value< 0.01 and p-value<0.001, respectively.

A normalized final score combining module score and negpro was used to represent the performance of a particular method on a given dataset. The score of csuWGCNA is significantly higher than both the signed and unsigned methods (BH adjusted P_csuWGCNA-signed_ =0.004, P_csuWGCNA-unsigned_ =0.003, Figure 4C). The unsigned method also performs better than the signed method (BH adjusted P value=0.003). This suggests that csuWGCNA can detect more negative correlations than signed and unsigned methods and performs as well as these two methods in detecting known modules.

### csuWGCNA captures more negatively correlated miRNA-targets and lncRNA-mRNA pairs

We next examined the improvement of csuWGCNA in complex human brain data. MiRNA and lncRNA are two types of non-coding RNA that have been reported to negatively regulate their target genes. We used two gene expression datasets that measured expression of miRNA and lncRNA from human brain, SMRI and BrainGVEX, as our test datasets.

We evaluated the methods from two aspects, the proportion of negative correlations by miRNA/lncRNA and the biological functions of the modules. The biological functions are represented by prediction of KEGG pathways. We tested 58,069 miRNA-target interactions (MTI) from miRTarBase in SMRI data and 7,334,095 lncRNA-gene pairs with significantly negative bicor values (FDR<0.05) in BrainGVEX data. Because the MTIs collected were validated experimentally, we only set the criteria that correlation<0 on them. The final score is a sum of z-scores of two metrics above.

Overall, the csuWGCNA performed best among three methods. At the gene level, csuWGCNA captured 16% of the MTIs and 33% of the lncRNA-gene pairs, which was more than that of signed and unsigned methods (chi-square test, P_csuWGCNA-unsigned_<1.89e-09, all other P<2.2e16, Figure 5A, Figure 5B). We found that 98% MTIs captured by csuWGCNA were functionally validated by RT-PCR, Western blot or RNA sequencing (Supplemental Figure 2). At the module level, we found that csuWGCNA and the signed method both detected the modules and have a lower FDR in prediction of KEGG pathway than the unsigned method (Figure 5C, Figure 5D), especially in the brainGVEX data with a large sample size. This suggested that csuWGCNA is able to detect modules corresponding to known biological pathways as well as the signed method and better than unsigned method. By combining these two criteria, final scores indicate that the csuWGCNA performs better than both signed and unsigned methods in SMRI and BrainGVEX data (Figure 5E, Figure 5F).

**Figure 5.**
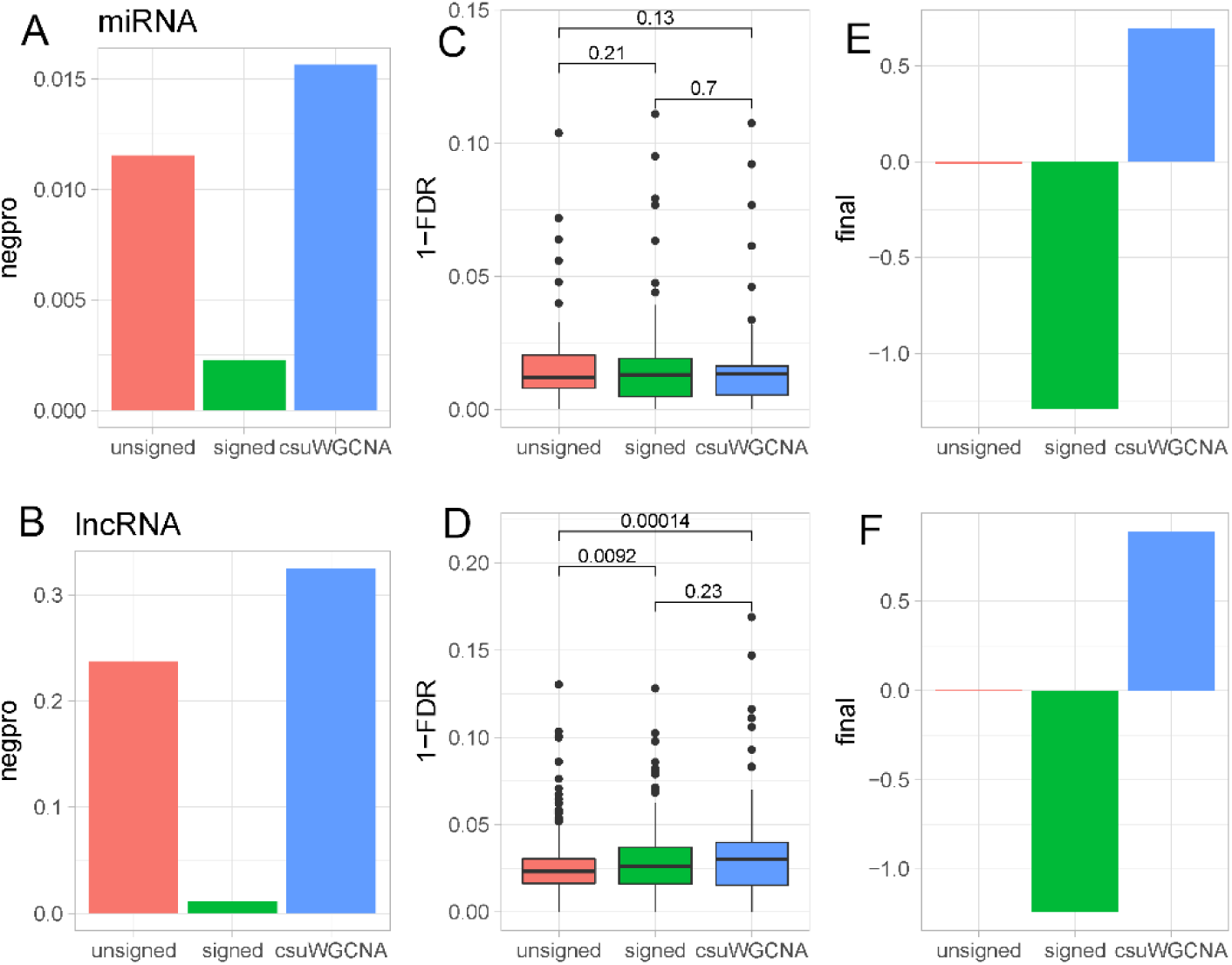
Summary of coexpression analysis of SMRI data (upper panel) and BrainGVEX data (bottom panel). (A-B) Negpro of miRNA - target and lncRNA - gene. (C-D) The FDR of predicting KEGG pathway (n=254, two-sample Wilcoxon test). (E-F) Final score of three method’s performance. The final score is a sum of zscore of detecting negtative correlations and predicting KEGG pathway.

## Discussion

Here we introduced csuWGCNA, which is a combination method of signed and unsigned WGCNA that captures more negative correlations. The csuWGCNA works by treating the positive and negative correlations equally and giving high connection strength to the weak and moderate correlations. We tested csuWGCNA on 22 datasets and compared the results to those of signed and unsigned methods. We showed that csuWGCNA is capable of detecting negative correlations and maintaining robust module detection in gene co-expression network analysis.

We showed the effectiveness of csuWGCNA from simple simulation data and complex human brain data. Our comparison includes, the detection of negative correlations and module reproducibility. In simulation data, we found that csuWGCNA was better than signed and unsigned methods in detecting weak and moderate correlations as predicted. This is an important improvement because most biologically negative correlations are small. We calculated the correlation between validated miRNA and targets and found that the majority of correlations are between −0.4~0 (Supplemental Figure 1). In ground truth data, we showed that csuWGCNA can reproduce known modules well in multiple datasets. The module score of csuWGCNA was not significantly improved. This was predicted because csuWGCNA aims to strengthen the detection of weakly correlated genes and global gene detection is still preserved in the three methods.

The real data is much more complex than the simulated data where gene co-expression is largely unknown. In the brain data, we showed that csuWGCNA can capture a higher proportion of negative MTIs and lncRNA-gene pairs than the other two methods. The importance of this is two-fold. First, we proved that the experimentally validated gene pairs have consistent co-expression at the whole transcriptome level in csuWGCNA. Second, it suggests promising application of csuWGCNA in detecting novel negative gene relationships. We also showed that csuWGCNA modules contain more functional pathways, especially compared to the unsigned method (p=0.13 in SMRI, p=0.0001 in BrainGVEX), even though the modules are preserved in both signed and unsigned networks (Supplemental Figure 3, Supplemental Figure 4). This suggests that csuWGCNA has the potential to obtain a larger number of informative modules. By extension, this further suggests that csuWGCNA may capture novel negative relationships related to specific biological functions.

The major advantage of csuWGCNA is that it is able to capture more disease-related negatively correlated lncRNA-mRNA pairs than the signed or unsigned methods. We found some lncRNAs and potential targets related to schizophrenia only by csuWGCNA. For example, lncRNA LINC00473 is a hub gene of a module downregulated in the schizophrenia (log2FC=-0.003, FDR=4.81E-05). LINC00473 is negatively correlated with TMEM245, which has been reported to be associated with negative symptoms of schizophrenia (P_meta-analysis_<6.22×10−6)^24^. This pair was only detected by the csuWGCNA method. As another example, NEAT1 is a brain-expressed lncRNA, which has been reported to change expressions in schizophrenia and neurodegenerative disease^25,26^. In the csuWGCNA module, NEAT1 negatively correlated with DPEP2 and ABCC12 which are downregulated in schizophrenia. In summary, csuWGCNA is a promising method to detect negatively correlated genes in transcriptome studies.

## Conclusion

csuWGCNA is an effective method for constructing gene co-expression networks. It can capture more negative correlations and maintain robust module detection compared to the original WGCNA procedure. Applying csuWGCNA on transcriptome data will help interpretations in systems biology.

## Methods

### Datasets and quality control

#### Simulation

We simulated 14 gene expression datasets containing 2000 genes and 50 samples using function simulateDatExpr. The proportion of negative correlations in each dataset was from 0.01 to 0.3, which was controlled by parameter propNegativeCor.

#### Ground truth data

We used six expression datasets for ground truth analyses, two from *E. coli*^27,28^, two from yeast (synapse.org/#!Synapse:syn2787209/wiki/70349), and two synthetic datasets. These datasets were collected by Saelens *et al*. in a comprehensive evaluation of module detection methods^1^ and were quality controlled and quantile normalized. The known modules were extracted from known gene regulatory networks^29–31^ used the strict module definition from Saelens *et al*. Strictly co-regulated modules were defined as groups of genes known to be regulated by exactly the same set of regulators. We downloaded the data from a Zenodo repository (doi: 10.5281/zenodo.1157938).

#### Human brain data

We used two different datasets from human brain: data from Stanley Medical Research Institute Neuropathology Consortium and Array collections (SMRI) and BrainGVEX, as real test data. The gene profiling and data pre-processing are described in Supplemental Materials.

##### SMRI data

We used parietal cortex tissue. The data measured the expression of miRNA and mRNA in SCZ and BD patients and controls. We removed non-Europeans, duplicates, and samples missing any of the mRNA or miRNA. After filtering, we retained 75 samples (25 SCZ, 25 BD, 24 controls), yielding data for 19,984 mRNAs and 470 miRNAs for subsequent analyses.

##### BrainGVEX data

We used RNA-Seq data of frontal cortex samples from the PsychENCODE project. The samples included 248 healthy control, 71 BD and 90 SCZ patient brains. Genes with Transcripts Per Million (TPM) lower than 0.1 in more than 25% of samples, mitochondrial genes, and pseudoautosomal genes were dropped. We calculated co-expression between samples, and samples with z-score normalized connectivity with other samples lower than −2 were removed. After filtering, 409 samples and 25774 genes were retained for subsequent analyses. Linear regression was used to remove the effect of covariates including age, sex, RIN, PMI, brain bank, batches, and principal components of sequencing statistics (seqPC). The seqPCs were the top 29 principal components of PCA on sequencing statistics. The covariates were selected by Multivariate adaptive regression splines (MARs).

### network construction

We completed signed, unsigned and csuWGCNA on all the datasets independently. Bicor was chosen to calculate the correlation between genes. We set power =12 for all the simulation datasets. Other parameters were as follows: ds = 2; minModSize = 30; dthresh = 0.1; pam = FALSE. For the ground truth data, the power of the three methods on each dataset calculated (Supplemental Table 1). Other parameters were as follows: ds = 4; minModSize = 20; dthresh = 0.2; pam = TRUE. For SMRI data, the soft power for signed, unsigned, and csuWGCNA was 5, 3 and 6, respectively. The parameters were as follows: TOMtype was unsigned, deepSplit was 4, minimum module size was 30 and mergeCutHeight wass 0.2, pamStage was true. For BrainGVEX data, the soft power for signed, unsigned, and csuWGCNA was 12, 4 and 10, respectively. The parameters were as follows: TOMtype was unsigned, deepSplit was 4, minimum module size was 40 and mergeCutHeight was 0.2, and pamStage was false. cutreeHybrid function was used to cut the gene tree.

### Evaluation metrics

We used three different types of metrics to evaluate signed, unsigned and WGCNA at gene pair and gene module levels. At the gene pair level, we used negpro, which is. At the gene module level, we considered the robustness and the biological functions of modules. The robustness of modules was used for comparison between known modules and observed modules in ground truth data. We used six classic metrics for comparing modules: specificity, sensitivity, NPV, PPV, relevance and recovery. Following are the formulas for the metrics where G represents all genes in a given dataset, M represents a gene set in an observed module, and m represents a gene set in a known module. For each M and m, the metrics are defined as:

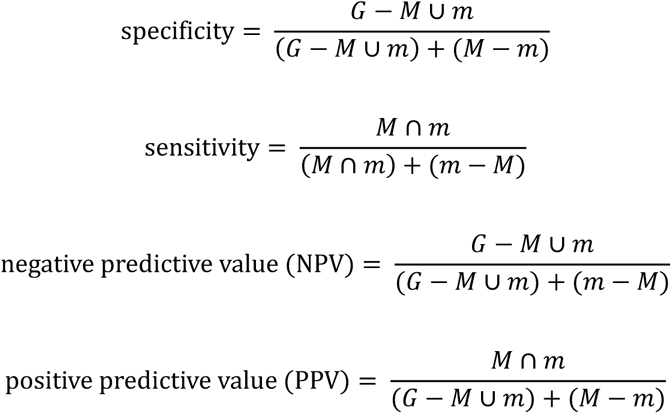

We compared every known module m and observed module M. For each M, we calculated the max value of a metric, such as specificity, sensitivity, or others, across all m. We averaged these max values to define the final value of a metric in a given dataset. The relevance and recovery are two metrics used to assess whether every observed module can be matched with a known module. We started by calculating the Jaccard index between every m and M.

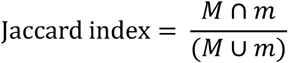

Relevance wass defined as the median value of the maximum Jaccard index value across all m for a given dataset. Recovery was defined as the median value of the maximum Jaccard index value across all M for a given dataset.

In the ground truth analysis, we calculated the module score as follows for evaluating module reproducibility.

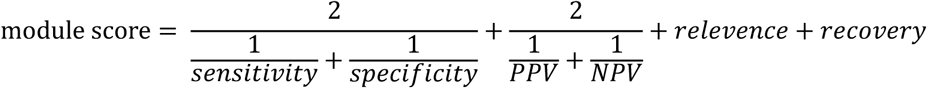

A final score combining module score and negpro was calculated.

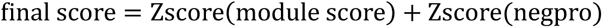

In the human brain data analysis, we replaced the module score with the prediction of from the KEGG pathway. The prediction of the KEGG pathway was represented by false discovery rate (FDR) defined as follows. For a KEGG pathway P and an observed module M, we calculated the minimum FDR across all M for a given P.

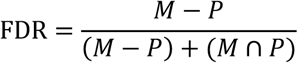

### Negative correlated miRNA-target interactions

To obtain the experimentally validated MTIs, we downloaded human MTIs from miRTarBase^32^. There were 502654 MTIs collected including 15064 genes and 2599 miRNAs. Among them, 117945 MTIs were detected by our SMRI data and 58069 MTIs showed negative correlations. The MTIs are cataloged by experimental evidence in miRTarBase. The strong evidence was considered to be reporter assay or Western blot and the weak evidence was considered to be microarray or pSILAC.

### KEGG pathways

To evaluate the prediction of biological pathway, 289 KGML files for human species were downloaded from the KEGG website^33^. The R package KEGGgraph^34^ was used to operate the KGML file and extract the gene members.

### Code Availability

The code of csuWGCNA is available from https://github.com/RujiaDai/csuWGCNA.

## Acknowledgement

This work was supported by NSFC grants 81401114, 31571312, the National Key Plan for Scientific Research and Development of China (2016YFC1306000), and Innovation-Driven Project of Central South University (No. 2015CXS034, 2018CX033) (to C.Chen), and NIH grants 1U01 MH103340-01, 1R01ES024988 (to C.Liu). All the data contributors are sincerely thanked for data provided.

